# Integrating Genomic and Transcriptomic Data to Reveal Genetic Mechanisms Underlying Piao Chicken Rumpless Trait

**DOI:** 10.1101/2020.03.05.978742

**Authors:** Yun-Mei Wang, Saber Khederzadeh, Shi-Rong Li, Newton Otieno Otecko, David M Irwin, Mukesh Thakur, Xiao-Die Ren, Ming-Shan Wang, Dong-Dong Wu, Ya-Ping Zhang

## Abstract

Piao chicken, a rare Chinese native poultry breed, lacks primary tail structures, such as pygostyle, caudal vertebra, uropygial gland and tail feathers. So far, the molecular mechanisms underlying tail absence in this breed have remained unclear. We employed comprehensive comparative transcriptomic and genomic analyses to unravel potential genetic underpinnings of rumplessness in the Piao chicken. Our results reveal many biological factors involved in tail development and several genomic regions under strong positive selection in this breed. These regions contain candidate genes associated with rumplessness, including *IRX4, IL-18, HSPB2*, and *CRYAB*. Retrieval of quantitative trait loci (QTL) and gene functions implied that rumplessness might be consciously or unconsciously selected along with the high-yield traits in Piao chicken. We hypothesize that strong selection pressures on regulatory elements might lead to gene activity changes in mesenchymal stem cells of the tail bud and eventually result in tail truncation by impeding differentiation and proliferation of the stem cells. Our study provides fundamental insights into early initiation and genetic bases of the rumpless phenotype in Piao chicken.

## Introduction

Body elongation along the anterior-posterior axis is a distinct phenomenon during vertebrate embryo development. Morphogenesis of caudal structures occurs during posterior axis elongation. The tail bud contributes most of the tail portion [1]. This structure represents remains of the primitive streak and Hensen’s node and comprises a dense mass of undifferentiated mesenchymal cells [1]. Improper patterning of the tail bud may give rise to a truncated or even absent tail [2]. Previous investigations implicated many factors in the formation of posterior structures. For example, loss of *T Brachyury Transcription Factor* (*T*) causes severe defects in mouse caudal structures, including the lack of notochord and allantois, abnormal somites, and a short tail [3]. Genetic mechanisms for rumplessness vary among the different breeds of rumpless chicken. For instance, Wang et al. revealed that rumplessness in Hongshan chicken, a Chinese indigenous breed, is a Z chromosome-linked dominant trait and may be associated with the region containing candidates like *LINGO2* and the pseudogene *LOC431648* [4,5]. In Araucana chicken, a Chilean rumpless breed, the rumpless phenotype is autosomal dominant and probably related to two proneural genes – *IRX1* and *IRX2* [6,7].

Piao chicken, a Chinese autochthonic rumpless breed, is native to Zhenyuan County, Puer City, Yunnan Province, China, and is mainly found in Zhenyuan and adjacent counties [8]. This breed has no pygostyle, caudal vertebra, uropygial gland and tail feathers [8], hence, an ideal model for studying tail development [9]. Through crossbreeding experiments and anatomical observations, Song et al. showed that rumplessness in Piao chicken is autosomal dominant and forms during the embryonic period even though no specific stage was identified [9,10]. However, until now, the genetic mechanisms of rumplessness in this breed have not yet been elucidated.

The advent of next-generation sequencing and microscopy has made it possible to probe embryonic morphogenesis through microscopic examination, to study phenotype evolution using comparative population genomics, and to assess transcriptional profiles associated with specific characteristics via comparative transcriptomics. In this study, we integrated these three methods to uncover the potential genetic bases of the rumpless phenotype in Piao chicken.

## Results

### Comparative genomic analysis identifies candidate regions for the rumpless trait in Piao chicken

To investigate the genetic mechanisms underlying rumplessness in Piao chicken, we employed comparative population genomics to evaluate population differentiation between the Piao chicken and control chickens with a normal tail. In total, we analyzed the genomes of 20 Piao and 98 control chickens, including 18 red junglefowls (RJFs), 79 other domestic chickens and one green junglefowl (GJF) as an outgroup (Table S1). We identified 27,557,576 single-nucleotide polymorphisms (SNPs), more than half of which (52.5%) were in intergenic regions, 42.1% in introns, and 1.5% in exons. Functional annotation by ANNOVAR [11] revealed that 292,570 SNPs (about 1%) caused synonymous substitutions, while 131,652 SNPs (approximately 0.5%) may alter protein structure and function through non-synonymous mutations or gain and loss of stop codons. A phylogenetic tree and principal component analysis (PCA) showed certain population differences between the Piao chicken and controls (**Figure 1**A–C). High genetic relatedness was found between Piao and domestic chickens from Gongshan, Yunnan (Figure 1C). One Piao chicken sample clearly separated from the other Piao chickens and was close to Chinese domestic chickens from Dali, Yunnan and Baise, Guangxi. Since Zhenyuan, Dali, and Baise are geographically close neighbors, it is possible that Piao chicken is beginning to mix with exotic breeds due to the emerging advancements in transport networks in these regions.

**Figure 1.**
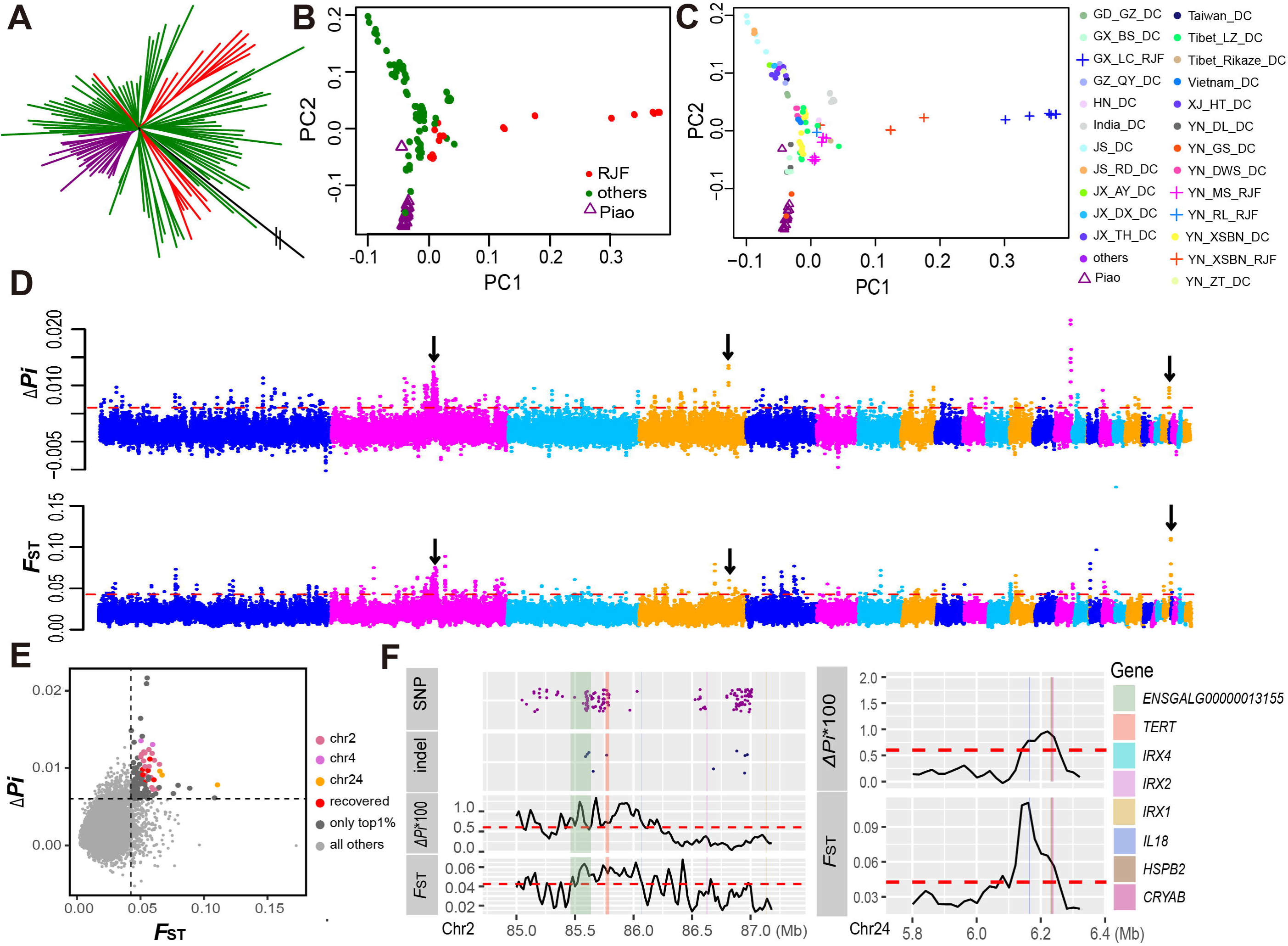
Population genomics analysis. **A**. Weighted phylogenetic tree of Piao chicken (purple) and controls grouped into RJF (red), GJF outgroup (black) and others (green). **B**. PCA plot of Piao and control chickens. Color groups are same as **A. C**. PCA plot similar to **B**, but marking control chickens from different places with different colors and shapes (cross is RJF, and the solid point represents other domestic chickens). GX_BS_DC, domestic chicken from Baise, Guangxi; YN_DL_DC, domestic chicken from DL, Yunnan; YN_GS_DC, domestic chicken from Gongshan, Yunnan; for the specification of other abbreviations please see Table S1. **D**. Manhattan plots of *F*_ST_ and Δ*Pi* based on the median sites of 40kb-sliding window regions. Black arrows point out strongly selected regions mentioned in the main text. **E**. Scatter plot for *F*_ST_ and Δ*Pi* of 40kb-sliding window regions. Larger dots are 112 selected sliding regions and colored if its recovery ratio of random sampling is greater than 0.95 in both *F*_ST_ and Δ*Pi*, while those in strongly selected regions mentioned in the main text are marked with different colors. **F**. Lines of *F*_ST_ and Δ*Pi*\*100 values plotted by the median sites of 40kb-sliding windows around the two strongly selected regions, i.e., chr2: 85.52–86.07Mb and chr24: 6.14–6.25Mb. Dark magenta and midnight blue dots represent highly differentiated SNPs and indels, respectively. The colored fillers are genes located in the two regions. Dashed lines indicate the top 1% threshold of descending *F*_ST_ and Δ*Pi*\*100.

Based on fixation index (*F*_ST_), nucleotide diversity (*Pi*) and genotype frequency (GF), we searched for genomic regions, SNPs, and insertions/deletions (indels) with high population differences between the Piao and control chickens (Materials and methods). We identified 112 40kb-sliding window genomic regions with strong signals of positive selection in the Piao chicken, out of which 28 were highly recovered in 1000 random samples of 20 controls from the 98 control chickens (Figure 1D–F and S1A, Table S2, and Materials and methods). Three regions with the strongest repeatable selection signals were chr2: 85.52–86.07Mb, chr4: 75.42–75.48Mb and chr24: 6.14–6.25Mb.

Chr2: 85.52–86.07Mb had a high proportion of highly differentiated (GF > 0.8) SNPs and indels, approximately 13.5% (66/488) and 10.4% (5/48), respectively (Figure 1F and Table S2). There are many important genes in this region. For instance, *IRX4*, along with *IRX1* and *IRX2*, belongs to the same *Iroquois* genomic cluster, *i*.*e*., the *IrxA* cluster [12]. Tena et al. discovered an evolutionarily conserved three-dimensional structure in the vertebrate *IrxA* cluster, which facilitates enhancer sharing and coregulation [12]. *IRX4* was reported to be involved in progenitor cell differentiation [13,14]. *Telomerase Reverse Transcriptase* (*TERT*) is implicated in spermatogenesis and male infertility [15]. In particular, *ENSGALG00000013155*, a novel gene, showed strong selection signals from both *F*_ST_ and *Pi*. This gene had 39 highly differentiated SNPs (23 located in introns and 16 in the intergenic region with *MOCOS*) and two highly differentiated indels (in introns). We retrieved expression profiles of this gene in different chicken tissues and development stages from three NCBI projects (SRA Accessions: ERP003988, SRP007412 and DRP000595) used in our previous study [16]. The results showed that the expression levels of this gene were high in three tissues, *i*.*e*., adipose, adrenal gland and cerebellum, and gradually increased during the early development of the chicken embryo (Figure S2A). The adrenal gland affects lipid metabolism and sexual behavior [17,18], while the cerebellum is involved in emotional processing [19]. These results suggest that *ENSGALG00000013155* plays important roles in fat deposition, sexual behavior and embryogenesis in chicken.

Chr4: 75.42–75.48Mb includes one gene – *Ligand Dependent Nuclear Receptor Corepressor Like* (*LCORL*). *LCORL* was reported as a candidate gene for chicken internal organ weight [20], horse body size [21], and cattle production performance [22].

Chr24: 6.14–6.25Mb exhibited the strongest selection signal on chromosome 24 and has a fundamental gene – *Interleukin 18* (*IL18*) (Figure 1F). *IL18* is well known as a proinflammatory and proatherogenic cytokine, as well as an IFN-γ inducing factor [23]. The gene modulates acute graft-versus-host disease by enhancing Fas-mediated apoptosis of donor T cells [24]. It can also induce endothelial cell apoptosis in myocardial inflammation and injury [25]. Interestingly, this gene has been reported to play prominent roles in osteoblastic and osteoclastic functions that are crucial for bone remodeling by balancing formation and resorption [26,27]. *IL-18* affects the bone anabolic actions of parathyroid hormone *in vivo* [26], and drives osteoclastogenesis by elevating the production of receptor activator of nuclear factor-κB (NF-κB) ligand in rheumatoid arthritis [27]. *IL-18* is also known as a trigger of NF-κB activation [25]. Sequestration of NF-κB in zebrafish can disrupt the notochord differentiation and bring about *no tail* (*ntl*)-like embryos [28]. This no tail phenotype can be rescued by the T-box gene *ntl* (*Brachyury* homologue) [28]. Indeed, in chicken, blocking NF-κB expression in limb buds gives rise to a dysmorphic apical ectodermal ridge, a decrease of limb size and outgrowth, as well as a failure of distal structures, through the interruption of mesenchymal-epithelial communication [29].

### Developmental transcriptome analysis reveals differentially expressed genes (DEGs) associated with tail development

The tail bud begins to form at the Hamburger-Hamilton stage 11 of chicken embryo development [30,31], and undergoes multidimensional morphogenesis in the subsequent three to ten days [32]. Thus, it should be possible to analyze transcriptional diversity associated with tail generation during this period. Based on this premise, we sought to outline the functional factors involved in caudal patterning using RNA-Sequencing data from 9 Piao and 12 control chicken embryos after seven to nine days of incubation (Table S1). To minimize bias, the control samples were collected from 6 Gushi and 6 Wuding chicken embryos (Figure S2B and Materials and methods).

An evaluation of gene expression identified 437 DEGs between the Piao and control chickens across the three developmental days (**Figure 2**A–B, Table S3, and Materials and methods), including the gene *T* that is implicated in tail development [3]. Gene ontology (GO) enrichment analysis of all DEGs by the database for annotation, visualization and integrated discovery (DAVID v6.8) [33] showed that many biological processes were related to posterior patterning, including muscle development, bone morphogenesis, somitogenesis, as well as cellular differentiation, proliferation and migration (Figure 2C).

**Figure 2.**
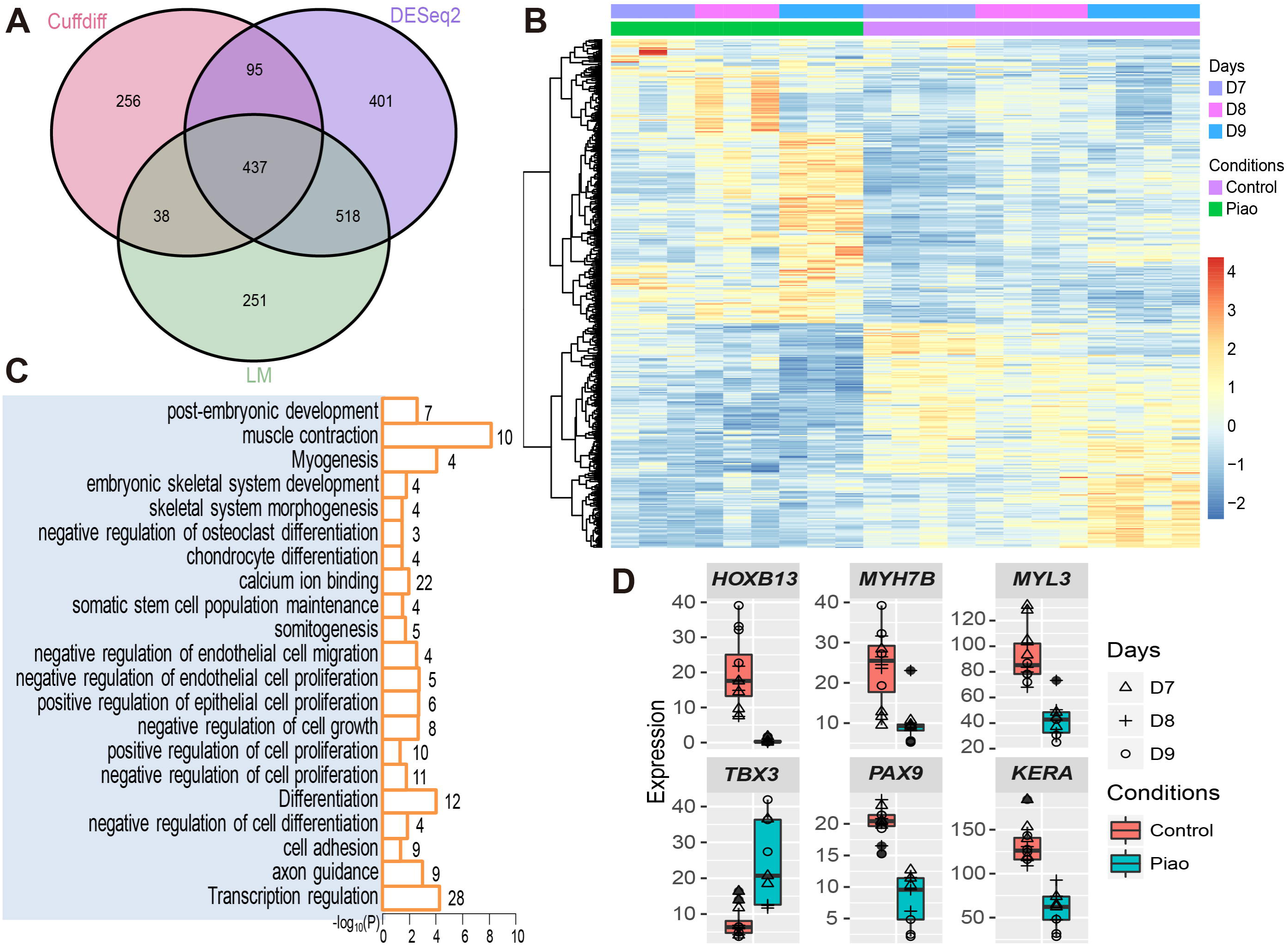
Differential expression analysis. **A**. Relationships of the number of DEGs from Cuffdiff, DESeq2, and the linear model (LM). **B**. Heatmap and hierarchical clustering dendrogram from FPKM of DEGs. Rows represent DEGs while columns show samples. **C**. Significant categories among DEGs by DAVID annotation. The number on the right of a bar presents gene number in the category. X-axis represents –Log_10_ *P* value. **D**. FPKM values of six representative DEGs in Piao and control chickens.

There were some interesting DEGs that had an expression fold change (FC) greater than 2 between the Piao and control chickens, including *T-box 3* (*TBX3*), *homeobox B13* (*HOXB13*), *myosin light chain 3* (*MYL3*), *myosin heavy chain 7B* (*MYH7B*), *KERA*, and *paired box 9* (*PAX9*) (Figure 2D). *TBX3*, encoding a transcriptional factor, had an expression in the Piao chickens three times higher than in controls. This gene regulates osteoblast proliferation and differentiation [34]. *HOXB13*, located at the 5’ end of the *HOXB* cluster, had an almost undetectable expression in the Piao chickens, but 36.7-fold higher levels in the controls. In mice, loss-of-function mutations in *HOXB13* cause overgrowth of the posterior structures derived from the tail bud, due to disturbances in proliferation inhibition and apoptosis activation [35]. *MYL3* showed 2.2 times lower expression in the Piao chickens compared to controls. This gene encodes a myosin alkali light chain in slow skeletal muscle fibers and modulates contractile velocity [36]. *MYH7B*, with 2.4-fold lower expression in the Piao chickens, encodes a third myosin heavy chain [37]. Mutations in *MYH7B* cause a classical phenotype of left ventricular non-compaction cardiomyopathy [37]. *KERA* encodes an extracellular matrix keratocan, which acts as an osteoblast marker regulating osteogenic differentiation [38]. The expression of *KERA* in the Piao chickens was 2.3-fold higher than in controls. *PAX9* and *PAX1* function redundantly to influence the vertebral column development [39]. Compared to controls, the expression level of *PAX9* in the Piao chickens is nearly 2.6 times lower. Analyzing gene expression patterns revealed that multiple genes are potentially involved in chicken tail development.

### Co-expression modules delineate the biological processes relevant to posterior patterning

To further elucidate principal biological pathways regarding caudal development, we constructed correlation networks through weighted gene co-expression network analysis (WGCNA) [40] (Materials and methods). We captured twelve co-expression modules, six of which (M4, M5, M7, M8, M9 and M10) were significantly correlated with the rumpless phenotype (Pearson correlation, *P* value < 0.05) (**Figure 3**A–G).

**Figure 3.**
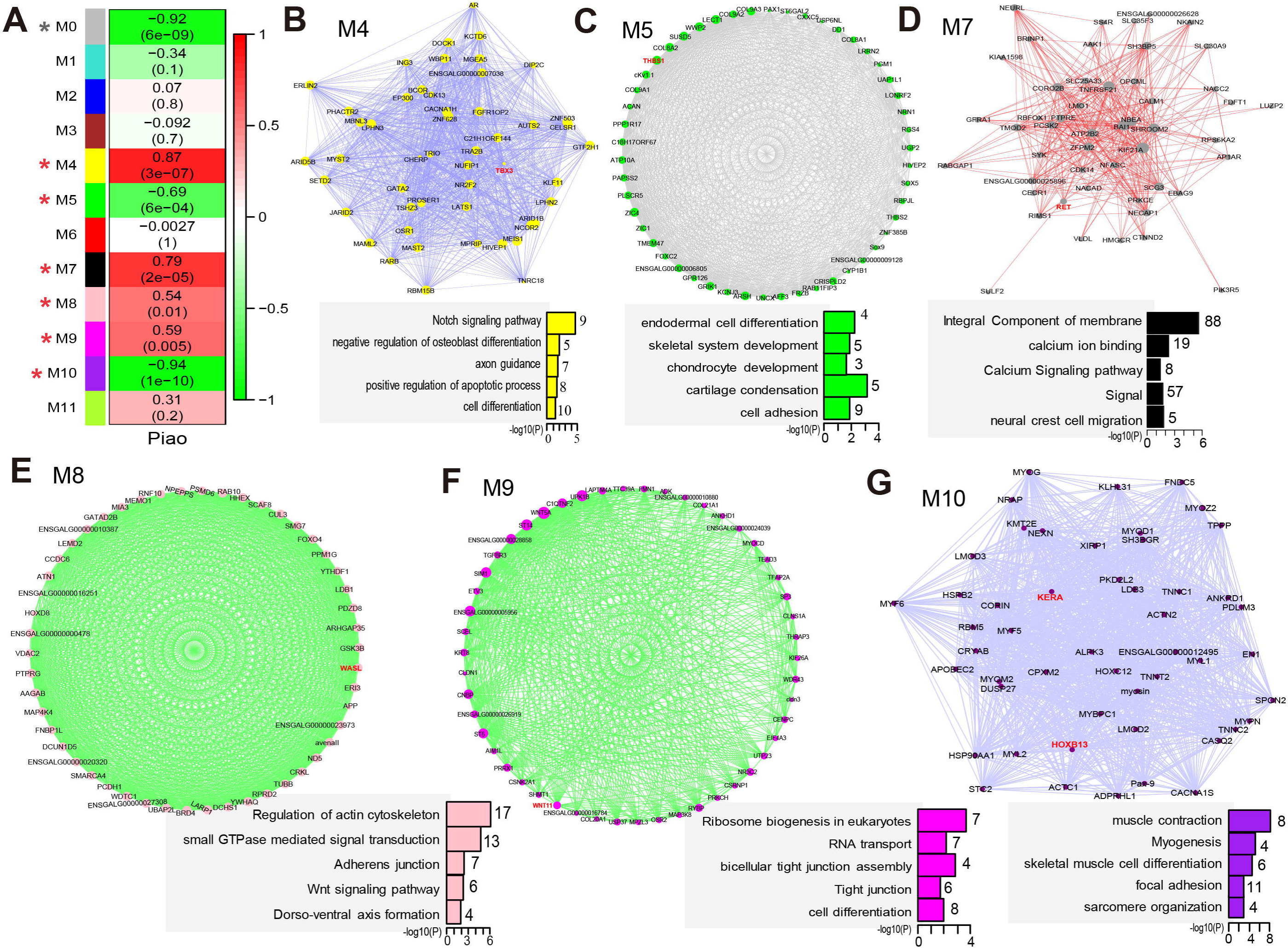
Gene co-expression module analysis. **A**. Module relationships with rumplessness. Values in and outside a parenthesis indicate *P* value and Pearson correlation coefficient, respectively. Modules were gradiently colored by Pearson correlation coefficients. Modules significantly correlated to rumplessness were marked with a red asterisk, while a grey asterisk labeled M0 to indicate that the module was excluded from the further analyses. **B–G**. Network plots from top 50 hub genes of six significant modules. Significant annotation categories of these modules from DAVID were colored according to network dots. Dot sizes indicate gene connectivity to others in the module.

Functional annotation for these significant modules revealed close linkages with embryonic development. Modules M5 and M10, which were negatively correlated with rumplessness, showed functional enrichment for skeletal system development and myogenesis, respectively (Figure 3C and G). Modules in positive correlation with rumplessness were implicated in several different pathways, including axon guidance and osteoblast differentiation for M4, calcium signaling pathway and neural crest cell migration for M7, actin cytoskeleton organization and dorso-ventral axis formation for M8, as well as transcription and tight junction for M9 (Figure 3B and D–F).

We searched for hub genes in each significant module and visualized the weighted networks with the top hubs (Figure 3B–G, Table S4, and Materials and methods). Interestingly, *TBX3*, one of the six DEGs mentioned above, was an M4 hub gene. Another five DEGs (*HOXB13, MYL3, MYH7B, KERA* and *PAX9*) are all M10 hubs. The M7 hub *ret proto-oncogene* (*RET*) encodes a transmembrane tyrosine kinase receptor with an extracellular cadherin domain [41]. *RET* induces enteric neuroblast apoptosis through caspases-mediated self-cleavage [42]. *Thrombospondin-1* (*THBS1*), an M5 hub gene, has effects on epithelial-to-mesenchymal transition and osteoporosis [43,44]. An M8 hub *WASL* is essential for Schwann cell cytoskeletal dynamics and myelination [45]. The M9 hub *WNT11* is crucial for gastrulation and axis formation [46,47]. These findings suggest that functional polygenic inter-linkages influence posterior patterning during chicken development.

### The only two DEGs under strong selection co-localized with *IL18*

To check whether our DEGs were strongly selected, we retrieved genes in the 40kb-sliding window regions with strong repeatable selection signals. We simultaneously searched for any highly differentiated SNP and indel in or flanking these DEGs. Finally, we only obtained two DEGs under strong selection, *i*.*e*., *Heat Shock Protein Beta-2* (*HSPB2*) and *Alpha-Crystallin B Chain* (*CRYAB*). Unfortunately, we found no highly differentiated SNP or indel related to these two DEGs. *HSPB2* and *CRYAB* are both located near *IL18* in chr24: 6.14–6.25Mb (Figure 1F). They encode a small heat shock protein and are essential for calcium uptake in myocyte mitochondria [48]. Nevertheless, they function non-redundantly: *HSPB2* balances energy as a binding partner of dystrophin myotonic protein kinase, while *CRYAB* is implicated in anti-apoptosis and cytoskeletal remodeling [49].

## Discussion

Accurate molecular regulation and control are vital for biological development and existence. Interfering these functional networks can lead to embryo death, diseases, deformities, or even the evolution of new characters [3–7]. Selection makes domestic animals achieve numerous phenotypic changes in morphology, physiology or behavior by modulating one or several components of primary biological networks [16].

In this study, we combined comparative transcriptomics and population genomics to explore the genetic mechanisms underlying rumplessness of Piao chicken. Our transcriptomic analyses presented many biological pathways that might be important to the late development of chicken tail. Genome-wide comparative analyses revealed several genomic regions under robust positive selection in the Piao chicken. These regions contain some fundamental genes, including *TERT, ENSGALG00000013155, IRX4, LCORL, IL-18, HSPB2*, and *CRYAB*, only two of which (*HSPB2* and *CRYAB*) were DEGs between the Piao and control chickens during D7–D9. Some of these genes might be associated with performance traits in the Piao chicken, such as *TERT* for egg fertilization, *ENSGALG00000013155* for fat deposition, and *LCORL* for body weight. Others might be implicated in axis elongation, such as *IRX4, IL-18, HSPB2* and *CRYAB*, through signaling pathways like NF-κB, calcium or apoptosis. Meanwhile, by retrieving all available QTL from the Animal QTLdb [50], we found that these regions were associated with production traits, such as growth, body weight, egg number, duration of broodiness and broody frequency (Table S2). In spite of lack of information about the evolutionary history of the Piao chicken, we know that this breed is characterized by high production, including elevated fat deposition rates, meat production, egg fertilization and egg hatchability [8,51]. Moreover, although Piao chicken originated in a relatively closed environment with limited genetic admixture with exotic breeds [51], it has high genetic variability and five maternal lineages [52], implying high hybrid fertility in the breed. Thus, we speculate that rumplessness, which exposes the posterior orifice of Piao chicken, might make intra-population or inter-population mating easier for the breed. This easier mating might increase egg fertilization, egg number, broody frequency and genetic variability in the Piao chicken. We propose that rumplessness might be consciously or unconsciously selected along with the high-yield traits of Piao chicken (**Figure 4**A).

**Figure 4.**
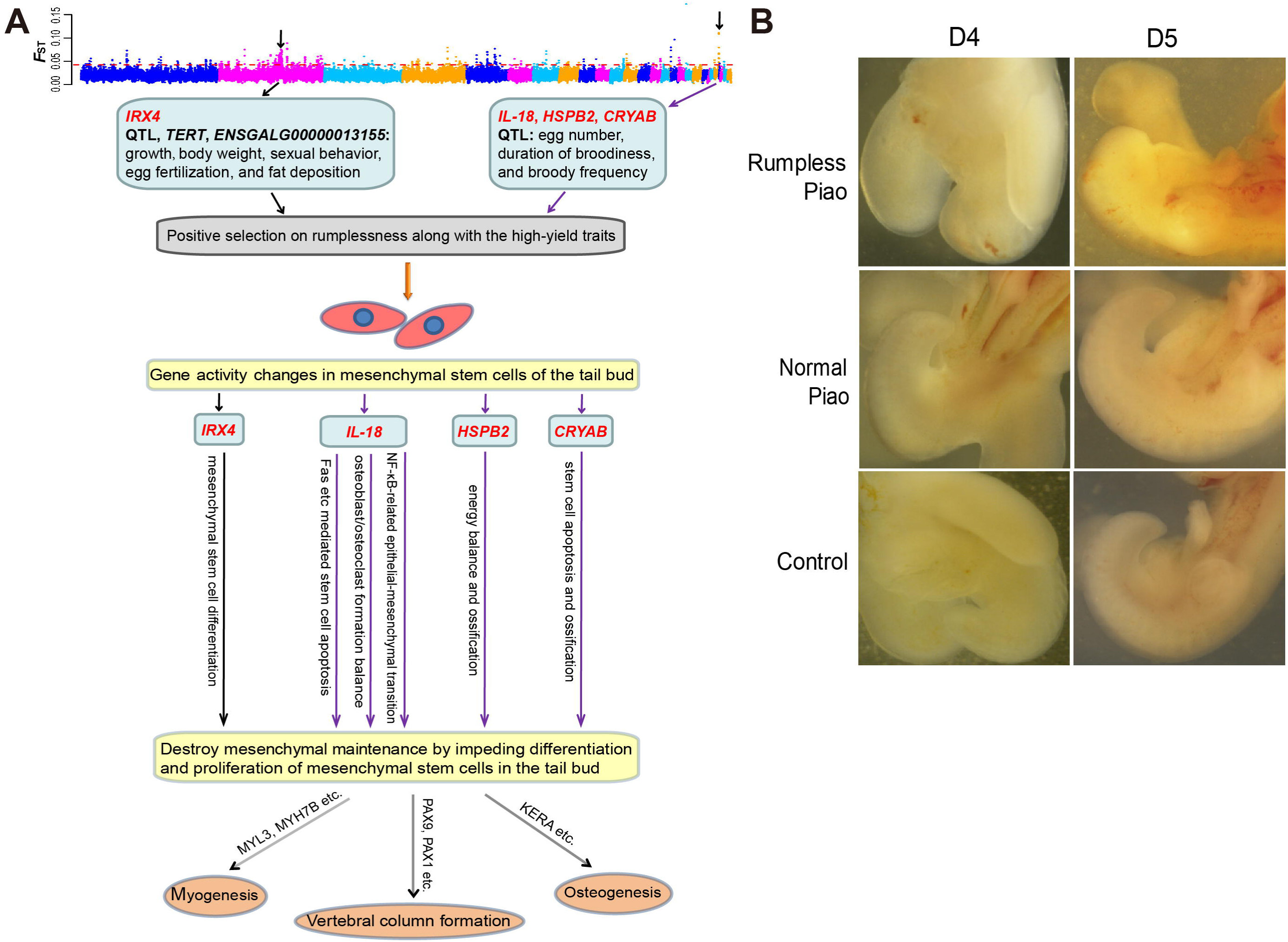
Proposed mechanisms underlying rumplessness in Piao chicken and microscopic examination of chicken tail embryogenesis. **A**. Rumplessness might be consciously or unconsciously selected along with the high-yield traits of Piao chicken. Strong positive selection pressures on regulatory elements of some candidate genes might lead to gene activity changes in the tail bud. These ectopic expressions would destroy mesenchymal maintenance by impeding differentiation and proliferation of mesenchymal stem cells in the tail bud, through multiple cell survival and differentiation pathways. This could then disrupt tail formation and prevent later developmental processes of the distal structures. **B**. The morphology of the posterior region during the fourth to fifth day of embryo development in rumpless Piao chicken, Piao chicken with a normal tail, and control chicken.

In addition, we found that all highly differentiated SNPs and indels, related to the above candidate genes, are located in noncoding regions or consist of synonymous mutations. Therefore, we hypothesize that positive selection pressures on the regulatory elements of some of these candidate genes would produce functional changes, which then leads to the rumpless phenotype in Piao chicken (Figure 4A). These activity changes likely take place at a very early development stage. In fact, when examining embryogenesis, we observed extreme tail truncation in most of the Piao embryos during the fourth day of incubation and later, while some Piao and all control embryos presented normal posterior structures (Figure 4B). The caudal morphogenesis in Piao embryos confirmed the descriptions from Song et al. [9] and Zwilling [53]. Song et al. found that the rumpless phenotype in Piao chicken is autosomal dominant [9]. Zwilling stated that dominant rumpless mutants arise at the end of the second day of embryo development and are established by the closure of the fourth day [53]. Previous studies have shown that the tail bud comprises a dense mass of undifferentiated mesenchymal cells and forms most of the tail portion [1]. The structure undergoes multidimensional morphogenesis from the third day of development [32]. Thus, we postulate that ectopic expressions, driven by positive selection pressures, might be initiated in the tail bud and impede normal differentiation and proliferation of the mesenchymal stem cells through signaling pathways like NF-κB, calcium or proapoptosis (Figure 4A). The result is the derailing of the mesenchymal maintenance in the tail bud and eventual failure of normal tail development. Hindrance to tail formation might have overarching impacts on later developmental processes of distal structures, for instance, *MYL3* and *MYH7B* involved in myogenesis, or *PAX9* and *PAX1* participating in vertebral column formation (Figure 4A). In particular, the strong selection on the proinflammatory cytokine gene *IL-18* could also be an adaptive response for a robust immunity, as rumplessness leaves the posterior orifice exposed to infections. However, as rumplessness in the Piao chicken is autosomal dominant, it is hard to confirm the rumpless phenotype and sample without RNA degradation before the fourth day of development. Therefore, we could barely even validate the expression of the identified genes in the tail bud further.

Previous studies have shown differences in the genetic architectures of taillessness among different chicken breeds [4–7], for example, Hongshan chicken [4,5] versus Araucana chicken [6,7]. Interestingly, Hongshan chicken has normal coccygeal vertebrae, while Araucana and Piao breeds have no caudal vertebrae and their rumplessness is autosomal dominant. Our results reveal potential similarities in the spatiotemporal bases of rumplessness in Piao and other dominant rumpless chickens [6,7,53]. This implies that autosomal-dominant rumplessness in chicken probably has the same genetic mechanisms and embryogenesis. By integrating comparative transcriptomics, population genomics and microscopic examination of embryogenesis, this study provides a basic understanding of genes and biological pathways that may be related, directly or indirectly, to rumplessness of the Piao chicken. Our work could shed light on the phenomenon of tail degeneration in vertebrates. Future endeavors should address the limitation to discern specific causative mutations that lead to tail absence in chicken.

## Conclusion

By combining comparative transcriptomics, population genomics and microscopic examination of embryogenesis, we reveal the potential genetic mechanisms of rumplessness in Piao chicken. This work could facilitate a deeper understanding of tail degeneration in vertebrates.

## Materials and methods

### Ethics statement

All animals were handled following the animal experimentation guidelines and regulations of the Kunming Institute of Zoology. This research was approved by the Institutional Animal Care and Use Committee of the Kunming Institute of Zoology.

### Whole-genome re-sequencing data preparation

DNA was extracted from blood samples from 20 Piao chickens by the conventional phenol-chloroform method. Quality checks and quantification were performed using agarose gel electrophoresis and NanoDrop 2000 spectrophotometer. Paired-end libraries were prepared by the NEBNext^®^ Ultra™ DNA Library Prep Kit for Illumina^®^ (NEB, USA) and then sequenced on the Illumina HiSeq2500 platform after quantification. 150bp paired-end reads were generated. Finally, we obtained tenfold average sequencing depth for each individual. Additionally, we used genomes from another 96 chickens with a normal tail from an unpublished project in our laboratory, including one outgroup GJF, 18 RJFs and 77 other domestic chickens (Table S1). Sequence coverage for these genomes ranged from about 5 to 110. Control genomic data from two additional individuals were downloaded from NCBI (SRA Accession: SRP022583) at https://www.ncbi.nlm.nih.gov/sra [54].

As the Piao chicken has been highly purified after a long time of conservation, it is hard to get enough normal-tailed Piao chicken samples with no inbreeding. Considering the short divergence time (about 8000 years) between domestic chickens and their ancestor – RJF [55], we used various normal-tailed chicken breeds as controls. Nevertheless, the number of each control breed was kept small. This design was based on previous studies [56,57]. In our opinion, the complex constitution of controls would reduce the background noises from some specific control breeds, but highlight the signatures of the target common trait, when compared with the Piao chicken where rumplessness is the main feature. Besides, comparing with exotic control breeds should weaken population differences among the Piao chicken, but highlight the common traits in this breed, like rumplessness.

### SNPs and indels detection

Variant calling followed a general BWA/GATK pipeline. Low-quality data were first trimmed using the software btrim [58]. Filtered reads were mapped to the galGal4 reference genome based on the BWA-MEM algorithm [59], with default settings and marking shorter split hits as secondary. We utilized the Picard toolkit (picard-tools-1.56, http://broadinstitute.github.io/picard/) to sort bam files and mark duplicates with the SortSam and MarkDuplicates tools, respectively. The Genome Analysis Toolkit (GATK, v2.6-4-g3e5ff60) [60] was employed for preprocessing and SNP/indel calling utilizing the tools RealignerTargetCreator, IndelRealigner, BaseRecalibrator, PrintReads, and UnifiedGenotyper. SNP variants were filtered using the VariantFiltration tool with the criteria: QUAL < 40.0, MQ < 25.0, MQ0 >= 4 && ((MQ0/(1.0*DP)) > 0.1). The same criteria were used for indels except setting: QD < 2.0 ‖ FS > 200.0 ‖ InbreedingCoeff < –0.8 ‖ ReadPosRankSum < –20.0.

### Population differentiation evaluation

To display the population structure of the Piao and control chickens, we built a phylogenic tree based on the weighted neighbor-joining method [61] for all SNP sites, and visualised it using the MEGA6 software [62]. We pruned SNPs based on linkage disequilibrium utilizing the PLINK tool (v1.90b3w) [63] with the options of ‘--indep-pairwise 50 10 0.2 --maf 0.05’. PCA was performed using the program GCTA (v1.25.2) [64] without including the GJF outgroup. Following the published formulas, we then computed *F*_ST_ [65] and *Pi* [66] values to detect signals of positive selection [56,66]. A 40kb sliding window analysis with steps of 20kb was performed for both *F*_ST_ and *Pi*. Nucleotide diversities (ΔPi) was calculated based on the formula ΔPi = *Pi* (Control) – *Pi* (Piao). The intersecting 40kb-sliding window regions in the top 1% of descending *F*_ST_ and Δ*Pi* were regarded as potentially selected candidates, an empirical threshold used previously [56]. Manhattan plots were drawn using the ‘gap’ package in R software [67] ignoring small chromosomes. To reduce false positive rates, we rechecked the candidate regions by analyzing 1000 replicates of 20 controls randomly sampled from all 98 control chickens. We identified intersecting 40kb-sliding window regions in the top 1% of descending *F*_ST_ and Δ*Pi* for each random sample. We then checked how many times the candidate regions were recovered by these intersecting regions during the 1000 random samples. We calculated the recovery ratio (i.e., times of recovery divided by 1000) for each candidate and defined those with a recovery ratio greater than 0.95 as the final strongly selected sliding regions.

GFs between the Piao and control chickens were compared to retrieve highly differentiated SNPs and indels. In general, there are three genotypes, *i*.*e*., 00, 01, and 11. 00 represents two alleles that are both the same as the reference genome. 01 represents one of the two alleles being altered, while the other is the same as the reference genome. 11 represents two alleles that are both altered. Considering sequencing errors, we defined a highly differentiated site using empirical thresholds: first, a site must exist in more than 15 (75%) Piao chickens and more than 50 (51%) control chickens; then, for the eligible site, sum of 01 and 11 must be larger than 0.8 in Piao chicken and less than 0.06 in control chickens. In total, we identified 488 and 48 highly differentiated SNPs and indels, respectively.

### Embryos microscopic observation

Fertilized Piao and normal-tailed control chicken eggs were purchased from the Zhenyuan conservation farm of Piao chicken and the Yunnan Agricultural University farm, respectively. Eggs were incubated at 37.5°C with 65% humidity. Embryo tail development was observed from the fourth to tenth day of incubation using a stereoscopic microscope.

### RNA isolation and sequencing

A total of 21 samples (9 Piao and 12 control chickens) were collected from the posterior end of embryos after seven to nine days of incubation and stored in RNA*later* at –80°C. RNA was extracted using Trizol reagent (Invitrogen) and RNeasy Mini Kits (Qiagen) and purified with magnetic oligo-dT beads for mRNA library construction. Paired-end libraries were prepared by the NEBNext^®^ Ultra™ RNA Library Prep Kit for Illumina^®^ (NEB, USA) and sequenced on the Illumina HiSeq2500 platform after quantification. 150bp paired-end reads were generated. Overall, we obtained approximately 5 Gbases (Gbp) of raw data for each library.

For the Piao chickens, there are three biological replicates for each of the three developmental days. By using samples from different developmental stages, we aimed to exclude inconsistent effects during development. Due to sampling difficulty, we used two Chinese native chicken breeds (6 Gushi and 6 Wuding chickens) as controls, in a way similar to the genomic analysis. Both control breeds have two biological replicates for each developmental stage. The Gushi and Wuding chicken are native to Henan and Yunnan, respectively, and both have a normal tail. In our opinion, using two control breeds would reduce trait noises from either breed, but strengthen signals from the common traits between the two breeds, such as having a normal tail compared with Piao chicken where rumplessness is the main feature.

### Transcriptomic data processing

We first trimmed low-quality sequence data using the software btrim [58]. Filtered reads were aligned to the chicken genome (galGal4.79, downloaded from http://asia.ensembl.org/index.html) using TopHat2 (v2.0.14) [68], with the parameters ‘--read-mismatches’, ‘--read-edit-dist’ and ‘--read-gap-length’ set to no more than three bases. We evaluated gene expression levels by applying HTSeq (v0.6.0) with the union exon model and the whole gene model, coupled with the Cufflinks program available in the Cufflinks tool suite (v2.2.1) [69] using default parameters.

### Correction and normalization

To improve the analyses, genes were filtered for expression in the three datasets: in at least 80 percent of the Piao or control samples, gene counts from both HTSeq models (union exon and whole gene) were no less than ten, while lower bound Fragments Per Kilobase of exon model per Million mapped fragments (FPKM) values from Cufflinks were greater than zero. We then performed normalization for gene length and GC content using the ‘cqn’ R package (v1.16.0) [70], based on the filtered count matrix of the HTSeq union exon model. The output values were defined as log_2_ (Normalized FPKM). The normalized matrix with genes kept in all three datasets was adjusted for unwanted biological and technical covariates, like development days, breeds, and sequencing lanes, via a linear mixed-effects model as previously described [71]. In detail, we calculated coefficients for these covariates with a linear model and then removed the variability contributing to them from the original log_2_ (Normalized FPKM) values. For example, when adjusting for development days, the number 1, 2 and 3 were used to replace D7, D8 and D9, respectively. We then calculated a coefficient for each gene using the “lm” function in R, with the number substitutes as a covariate. We removed the product of the coefficient and the number substitute from the log_2_ (Normalized FPKM) value to obtain the adjusted value. The adjusted data was then used for co-expression network construction.

### Differential expression analysis

We applied three methods to identify DEGs. First, 826 DEGs (FDR < 0.05) were identified by the Cuffdiff program in the Cufflinks tool suite with default parameters, using the bam files from TopHat2. Second, 1451 DEGs (FDR < 0.05) were found by DESeq2 [72] based on the read count data from the HTSeq union exon model. Third, 1244 DEGs (FDR < 0.05) were obtained with a linear model, where we used the log_2_ (Normalized FPKM) matrix from the normalization step and treated development stages, chicken breeds and sequencing lanes as covariates. In total, 437 DEGs found by all three methods were used as the final DEGs.

### Gene co-expression network analysis

To unravel underlying functional processes and genes associated with tail development, we carried out WGCNA in the R package [40] with a one-step automatic and ‘signed’ network type. The soft thresholding power option was set to 12 based on the scale-free topology model, where topology fit index R^^^2 was first greater than 0.8. The minimum module size was limited to 30. A height cut of 0.25 was chosen to merge highly co-expressed modules (*i*.*e*., correlation greater than 0.75). Finally, we obtained a total of 12 modules. M0 consisted of genes that were not included in any other modules, and thus was excluded from further analyses. We performed Pearson correlation to assess module relationships to the rumpless trait, and defined *P* value < 0.05 as a significant threshold.

### Hub genes and network visualization

In general, genes, which have significant correlations to others and the targeted trait, are the most biologically meaningful and thus defined as ‘hub genes’. Here, we referred to module eigengenes (MEs) as ‘hub genes’ dependent on high intramodular connectivity, and absolute values of gene significance (GS) and module membership (kME) greater than 0.2 and 0.8, respectively. GS values reflect tight connections between genes and the targeted trait, while kME mirrors eigengene-based connectivity between a gene expression profile and ME, and is also known as module membership [40]. The intramodular connectivity values measure the co-expression degree of a gene to other MEs in the module where it belongs. To visualize a weighted network, we ranked hub genes for each module by intramodular connectivity in descending order, and exported network connections between the top 50 hubs into an edge file with a topological overlap threshold of 0.1. The edge files were input to Cytoscape [73] for network analysis. Network plots for modules significantly related to the rumpless trait were then displayed based on decreasing degree values.

### Data access

Raw sequence data for the RNA samples and DNA samples of Piao chicken reported in this paper were deposited in the Genome Sequence Archive [74] in BIG Data Center [75], Beijing Institute of Genomics (BIG), Chinese Academy of Sciences (GSA Accession: CRA001387) at http://bigd.big.ac.cn/gsa. Codes and input files for the major analytic processes were stored in GitHub (accession is the title of this paper) at https://github.com.

## Authors’ contributions

YPZ, DDW and MSW designed the study. YMW and SK performed data analyses. YMW and SRL finished embryo incubation and observation. YMW and DDW wrote the manuscript. SK, NOO, DMI and MT revised the manuscript. MSW helped with comparative genomic analyses. XDR submitted the data. All authors read and approved the final manuscript.

## Acknowledgments

This work was supported by the Bureau of Science and Technology of Yunnan Province (2015FA026), the Youth Innovation Promotion Association, Chinese Academy of Sciences, and the National Natural Science Foundation of China (grant numbers 31771415, 31801054). We are grateful to the team of Prof. Yong-Wang Miao, Yunnan Agricultural University, for blood collection from adult Piao chickens. We also thank the support of the CAS-TWAS President’s Fellowship Program for Doctoral Candidates.

## Supplementary materials

**Figure S1 Random sampling recovery ratio of the 112 strongly selected 40kb-sliding window regions**

The top two bar plots show the recovery ratio for *F*_ST_ and Δ*Pi* in 1000 random samples of 20 controls from the 98 control chickens. The blue dashed lines indicate a recovery ratio of 0.95. Red asterisks show sliding window regions with a recovery ratio greater than 0.95 in both *F*_ST_ and Δ*Pi*, while the short black lines above the asterisks indicate candidate genes in the sliding regions. The bottom two line charts display *F*_ST_ and Δ*Pi* values based on the 98 control samples. Dashed lines indicate the top 1% threshold of descending *F*_ST_ and Δ*Pi*. The colored fillers present sliding window regions in the selected regions mentioned in the main text. The 40kb-sliding window regions are presented based on their median sites in order of chromosomal position.

**Figure S2 Expression levels of *ENSGALG00000013155* and chicken pictures**

**A**. FPKM values of *ENSGALG00000013155* in different chicken tissues and development stages from the three NCBI projects (SRA Accessions: ERP003988, SRP007412 and DRP000595) that were used in our previous work [16]. **B**. The pictures of male (M) and female (F) individuals of Piao chicken, Gushi chicken, and Wuding chicken.

**Table S1 RNA and DNA sample information**

**Table S2 Strongly selected regions, highly differentiated SNPs and indels, as well as QTL annotations**

**Table S3 Expression values of DEGs and their DAVID annotation**

**Table S4 Hub genes of modules significantly correlated to rumplessness**

