## Supplementary figures and images for "Integrating Genomic and Transcriptomic Data to Reveal Genetic Mechanisms Underlying Piao Chicken Rumpless Trait"

### Figure S1

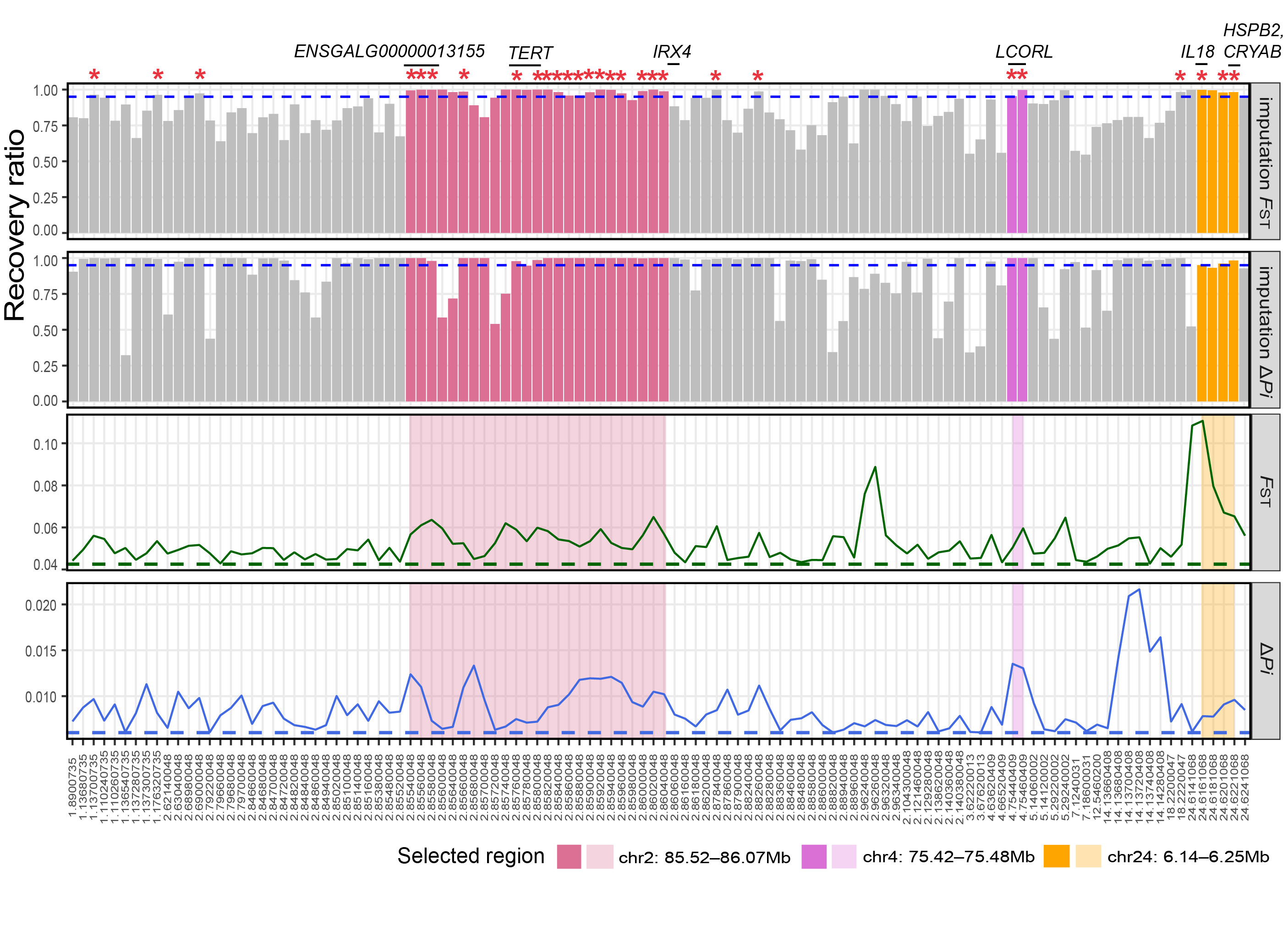

### Figure S2

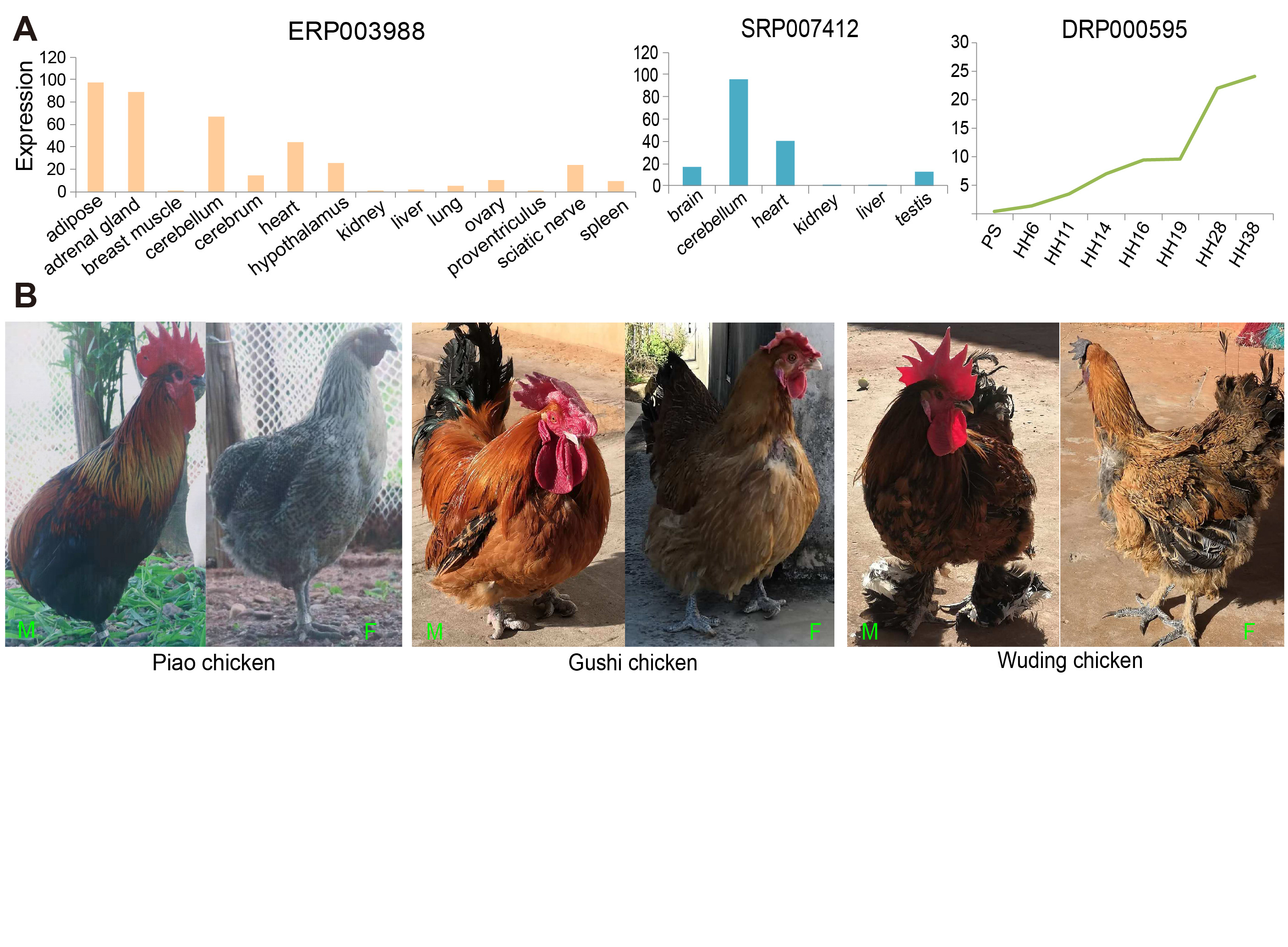
